# N1-methyladenine safeguards against aberrant RNA-protein association during proteostasis stress

**DOI:** 10.1101/667543

**Authors:** Marion Alriquet, Giulia Calloni, Adrían Martínez-Limón, Riccardo Delli Ponti, Gerd Hanspach, Martin Hengesbach, Gian G. Tartaglia, R. Martin Vabulas

**Affiliations:** Buchmann Institute for Molecular Life Sciences, Goethe University Frankfurt, Frankfurt am Main, Germany; Institute of Biophysical Chemistry, Goethe University Frankfurt, Frankfurt am Main, Germany; Centre for Genomic Regulation (CRG), The Barcelona Institute for Science and Technology, Barcelona, Spain; ICREA and Universitat Pompeu Fabra (UPF), Barcelona, Spain; Institute for Organic Chemistry and Chemical Biology, Goethe University Frankfurt, Frankfurt am Main, Germany; Department of Biology ‘Charles Darwin’, Sapienza University of Rome, Rome, Italy; Department of Neuroscience and Brain Technologies, Istituto Italiano di Tecnologia, Genoa, Italy

## Abstract

Post-transcriptional modifications of nucleotide bases can effect several aspects of mRNA function. For example, the recent work has established the role of m^6^A in the coordinated regulation of transcriptome turnover and translation during cellular differentiation and tumorigenesis. The levels of m^1^A in mRNAs reach up to 10% of those of m^6^A, yet the functional consequences of this modification are much less clear. Here we show that N1-methyladenine protects mRNAs against aberrant interactions during heat shock and amyloidogenesis in mammalian cells. The m^1^A methylation motif correlated with the enhanced sequestration of transcripts in stress granules (SG). The cognate methyltransferase TRMT6/61A accumulated and m^1^A was enriched in SG. Downregulation of the catalytic subunit TRMT61A enhanced amyloidogenesis in the cytosol and increased bystander protein and RNA co-aggregation with Aβ aggregates. Faulty granulation of mutant RNAs has been implicated in pathogenesis of protein aggregation disorders. Our results demonstrate that also normal mRNAs succumb to co-aggregation with proteins if RNA dynamics during stress is disturbed due to the insufficient N1-adenine methylation.

---

The target motif for the N1-adenine methylation on mRNAs by TRMT6/61 has been identified (*1, 2*). We analyzed its presence in SG-sequestered mRNAs (*3*) using a motif which can form an at least two base pair-long stem (Fig. 1a). The analysis revealed that TRMT6/61A-targeted transcripts were enriched in SGs (Fig. 1b). Two GO categories in SG-sequestered mRNAs with the m1A motif were significantly increased: “Regulation of RNA metabolic processes” (1.7x, 59 proteins) and “Axonogenesis” (4.6x, 13 proteins). The latter category represents an intriguing hit (Fig. S1a) because the build-up of the long process of a neuron requires packing and transport of RNA granules to the sites of local translation (*4*). The enrichment of axonogenesis proteins suggests that the m^1^A-related granulation is not restricted to stress conditions but might be a more general mechanism of the mRNA metabolism. Structurally, the m^1^A-motif containing SG-enriched transcripts were longer than control mRNAs (Fig. S1b).

**Figure 1.**
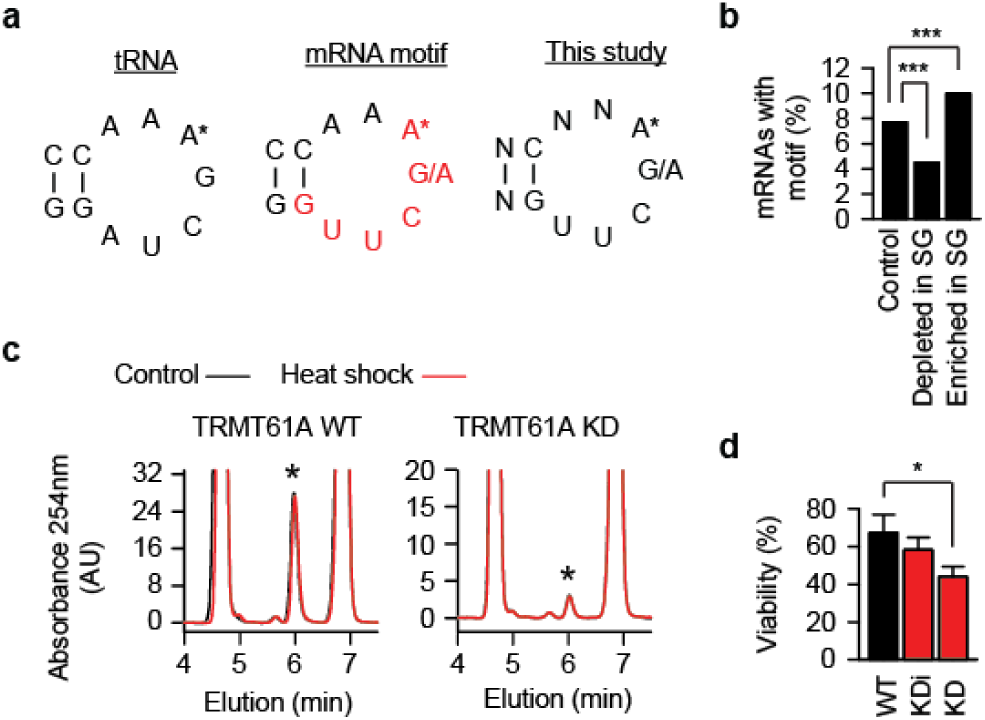
RNA methyltransferase TRMT6/61A is required during acute proteostasis stress. **a**, tRNA, TΨC arm of tRNA where adenine 58 (A*) is N1-methylated by TRMT6/61A. mRNA, mRNA motif (red) targeted by TRMT6/61A for adenine N1-methylation (*2*). This study, motif used here to predict mRNA targets of TRMT6/61A-mediated adenine N1-methylation. **b**, Fraction of mRNAs enriched or depleted in stress granules (SG) (*3*) and containing the m^1^A motif. Control, mRNA set neither enriched nor depleted in SG, ***p<0.001, chi-square analysis. **c**, HPLC analysis of N1-methylated adenosine (*) from cellular tRNAs under normal temperature (37°C) and after 1 h at 45°C. **d**, Reduced amounts of the methyltransferase correlate with the increased sensitivity to heat shock. WT, wild-type cells with normal amount of TRMT61A; KD and KDi, knock-down cells with strongly reduced and intermediate amounts of TRMT61A, respectively. *p<0.05, two-tailed t-test; N=3 independent experiments (mean + SD).

Next, TRMT61A-deficient cells were prepared and named “knock-downs” (KD) because of the residual levels of the catalytic subunit (Fig. S1c). Protein synthesis and cellular viability were not significantly changed and the proliferation was only slightly slower in KD cells (Fig. S1d-f). Heat shock for 60 min did not change the amount of m^1^A in tRNAs (Fig. 1c and Fig. S1g), yet affected cellular viability correlating with TRMT61A levels (Fig. 1d). Similarly, TRMT61A activity was needed to protect cells from another acute proteostasis stressor, arsenite (Fig. S1h, i).

N1-adenine methylation on mRNAs is known to increase upon heat shock (*5*), which suggests the link between m^1^A in mRNAs and the impaired survival of KD cells. In support, TRMT6/61A methyltransferase localized to SG under stress (Fig. 2a and Fig. S2a). To quantify N1-methyladenine accumulation in SG by mass spectrometry, we established SG isolation (*6*) and targeted selected ion monitoring procedures (Fig. S2b-d). A significant enrichment of m^1^A (calculated as m^1^A/A fraction) in SG was detected as compared to cytosolic mRNAs (Fig. 2b). The origin of the signal from ribosomes in SG can be excluded, because the only m^1^A on human ribosome is on the 60S subunit (*7*), which is excluded from SG (*8*).

**Figure 2.**
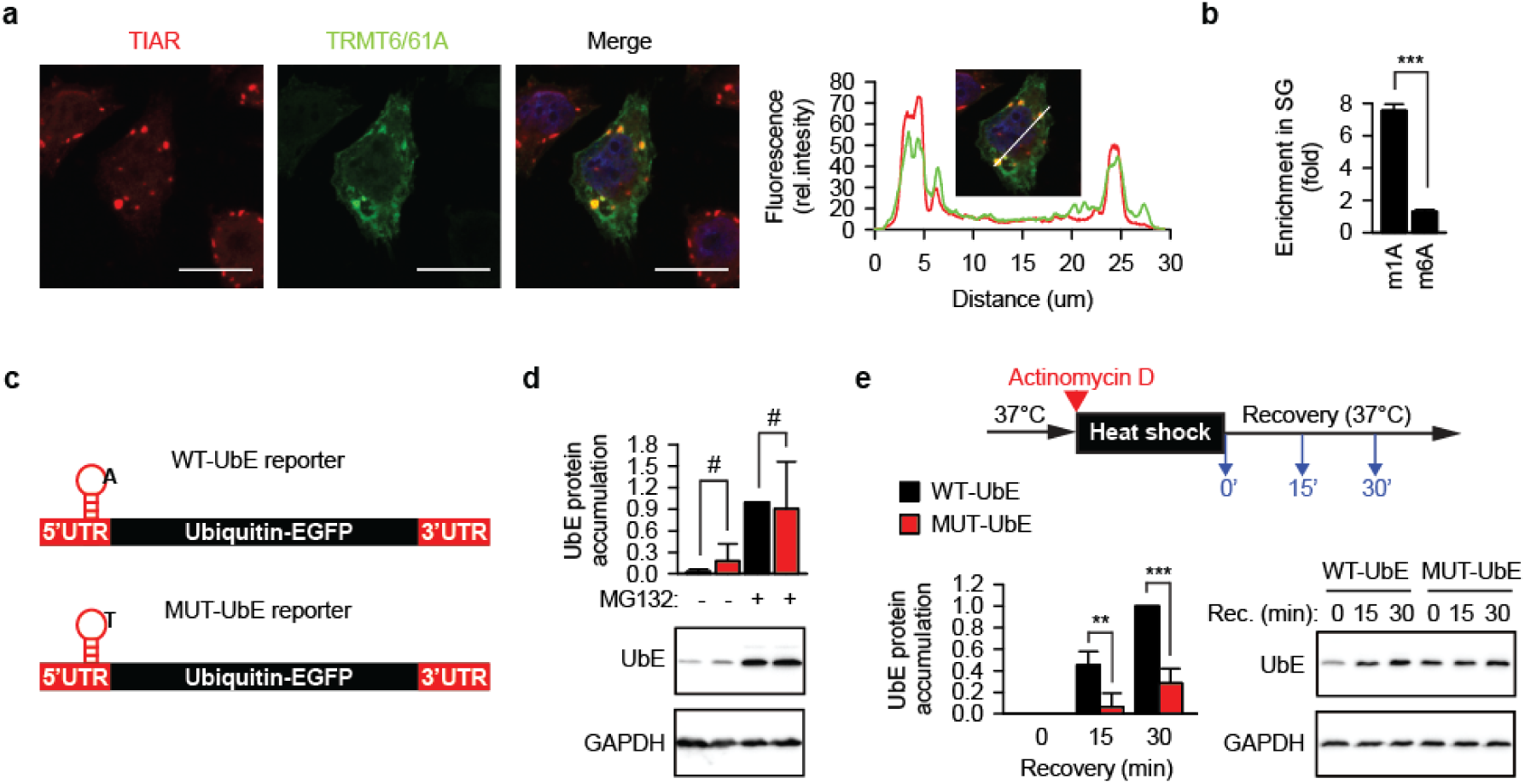
TRMT6/61A safeguards mRNA during heat shock. **a**, Localization of TRMT6/61A in arsenite-induced stress granules that are detected with anti-TIAR antibody. A representative image from three independent experiments. Scale bar 20 μm. The bar in the inset picture indicates the line where the fluorescence profiles were extracted. Red, TIAR; green, TRMT6/61A. **b**, N1-methyladenosine is enriched in stress granules compared to cytosolic polyA mRNAs as measured by mass spectrometry. m1A, fraction of N1-methyladenosine over unmodified adenosine; m6A, fraction of N6-methyladenosine over unmodified adenosine. The respective fraction in cytosolic polyA mRNAs was set as 1. ***p<0.001, two-tailed t-test; N=3 independent experiments (mean + SD). **c**, Schematic depiction of the m^1^A motif-containing reporter. UTR, untranslated region. The position of the motif on the transcript is indicated. The nucleotide sequence of the motif is shown in Figure S5G. **d**, Accumulation of the Ubiquitin-EGFP protein (UbE) from WT-UbE and MUT-UbE transcripts during 3 h-long inhibition of proteasomal degradation with 20 μM MG132 (+MG132). One representative anti-EGFP western blot from three independent experiments is shown. GAPDH was used as loading control. #, not significant difference; two-tailed t-test; N=4 independent experiments (mean + SD). **e**, Accumulation of UbE protein from WT-UbE and MUT-UbE transcripts during recovery after heat shock in the presence of actinomycin D to block synthesis of new mRNA molecules. UbE amount at timepoint 0 was subtracted from all values during recovery. One representative anti-EGFP western blot is shown. GAPDH was used as loading control. ***p<0.001, **p<0.01, two-tailed t-test; N=4 independent experiments (mean + SD).

To test functionally whether TRMT6/61A can protect mRNAs during proteostasis stress, m^1^A motif-containing reporters were generated and analyzed. An m^1^A motif from the 5’-UTR of the PRUNE1 transcript (*1, 2*) was inserted into the 5’-UTR of an Ubiquitin-EGFP (UbE) construct (Fig. 2c, Fig. S2e). The UbE protein is highly unstable and its accumulation can be detected with the precision of several minutes upon inhibition of proteasomal degradation in HeLa cells (*9*). Consequently, the accumulation of the UbE protein would indicate the functional amount of its coding mRNA in the cytosol at the time-point when degradation and transcription are inhibited. In parallel to the wild-type reporter WT-UbE, a control reporter MUT-UbE with mutated adenine was generated and tested under normal conditions (Fig. 2d). To test the reporters during heat shock and recovery, transcription of new mRNA was inhibited (Fig. S2f). Notably, heat-misfolded proteins overloaded the proteasomal capacity leading to small accumulation of protein from MUT-UbE (Fig. 2e, lane 4), which suggested that heat-related inactivation of MUT-UbE mRNA was less efficient. Oppositely, upon returning to 37°C, the protein from WT-UbE accumulated significantly more (Fig. 2e). These data indicate that the m1A motif-containing mRNA 1) during heat shock are sequestered more efficiently and 2) during recovery became functional again faster.

We analyzed whether the impaired dynamics of mRNA in cytosol can aggravate protein aggregation. The amyloid-β peptide Aβ_1-42_ fused to GFP (Aβ-GFP) was used to this end. Western blot analysis revealed an enhanced accumulation of Aβ-GFP in KD cells (Fig. 3a). Fluorescence microscopy confirmed the reduced capacity of the TRMT61A-deficient cells to constrain amyloidogenesis (Fig. 3b and Fig. S3a). The reintroduction of TRMT6/61A activity into KD cells significantly suppressed aggregation (Fig. 3c). Enzymatically impaired TRMT61A mutant D181A was less efficient in this suppression (Fig. S3b), which argues for the necessity of enzymatic function for mRNA safeguarding. We wondered if the m^1^A tag can rescue its containing transcripts also under chronic proteostasis stress imposed by amyloidogenesis. Reporters encoding NAD(P)H:quinone oxidoreductase 1 (NQO1), a stable human protein, were prepared (Fig. 3d and Fig. S3c). Transient co-expression of WT-NQO1 with Aβ-GFP resulted in reduced accumulation of the reporter protein (Fig. 3e). However, this effect was significantly stronger in the case of MUT-NQO1. The amounts of both reporter mRNAs were similar as confirmed by quantitative PCR (Fig. S3d), which suggests that the methylation of adenine in the TRMT6/61A motif was needed to sustain the functionality of the reporter mRNA in the presence of amyloid. On the other side, accumulation of NQO1 from wild-type and mutant reporters was similar when compared in KD cells supporting the centrality of the TRMT6/61A-dependent mechanism (Fig. S3e). The difference between reporter functionality in the presence of Aβ-GFP could be reproduced in another cell line, the murine melanoma B16-F10 (Fig. S3f).

**Figure 3.**
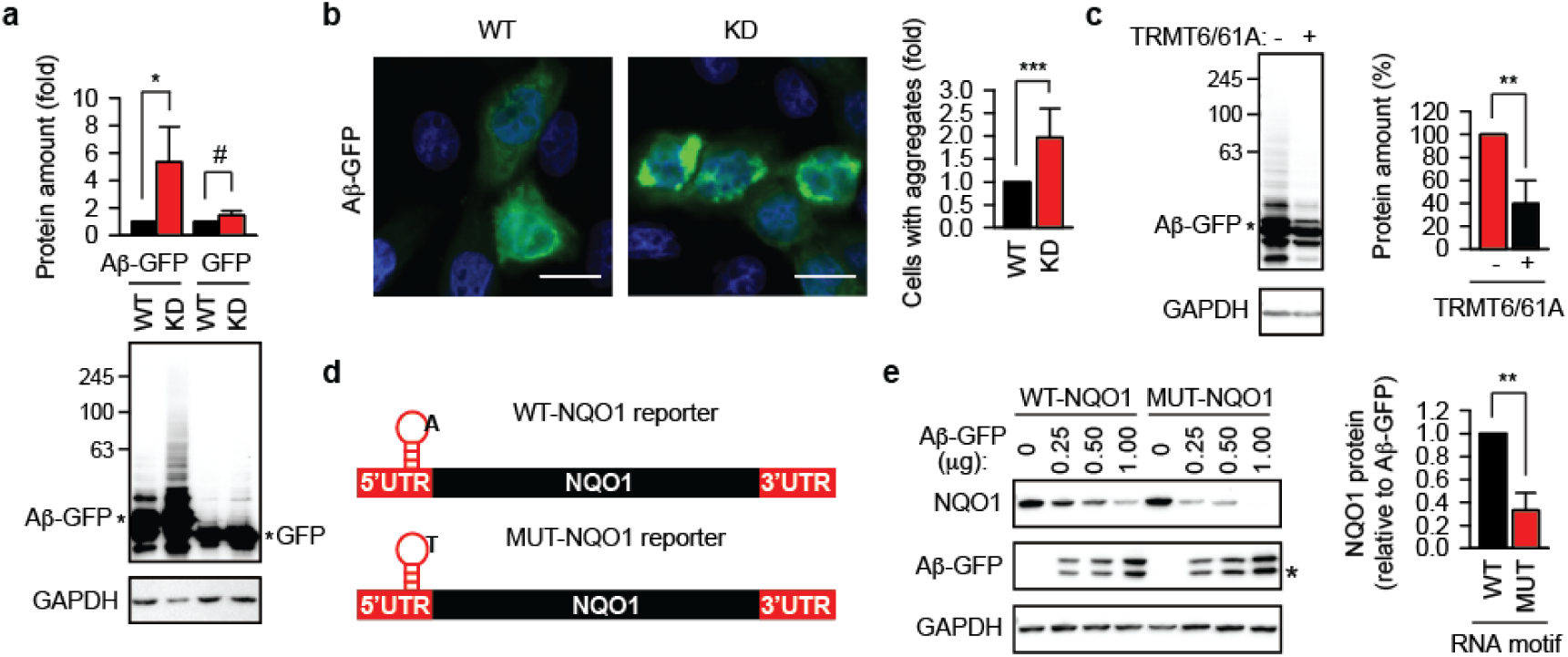
Lack of the TRMT6/61A-dependent mRNA methylation enhances amyloidogenesis. **a**, Western blotting of overexpressed Aβ-GFP and GFP. GAPDH was used as loading control. *p<0.05; #, not significant difference; two-tailed t-test; N=3 independent experiments (mean + SD). **b**, Aggregation of Aβ peptide-GFP fusion protein (Aβ-GFP, green) transiently overexpressed in wild-type (WT) and TRMT61A knock-down (KD) cells. Scale bar 20 μm. ***p<0.001, chi-square test on cumulative data from three independent experiments (N=3, mean + SD). **c**, Western blotting of overexpressed Aβ-GFP in TRMT61A knock-down cells. TRMT6 and TRMT61A were cotransfected as indicated. GAPDH was used as loading control. **p<0.01, two-tailed t-test; N=3 independent experiments (mean + SD). **d**, Schematic depiction of the m^1^A motif-containing NQO1 reporters. UTR, untranslated region. The position of the motif on the transcript is indicated. The nucleotide sequence of the motif is shown in Figure S6B. **e**, Accumulation of NQO1 protein from WT-NQO1 and MUT-NQO1 transcripts 24 h after cotransfection with indicated amounts of Aβ-GFP. One representative anti-FLAG western blot (NQO1) is shown. GAPDH was used as loading control. For quantitative comparison, NQO1 amount per unit of Aβ-GFP was calculated and averaged for all three Aβ-GFP concentrations. This value for WT-NQO1 reporter was set as 1. **p<0.01, two-tailed t-test; N=4 independent experiments (mean + SD).

Aberrant protein-protein interactions are known to be important for proteotoxicity (*10, 11*). Mechanistically, we entertained the possibility that mislocalized mRNAs become entrapped in protein aggregates and enhance protein co-aggregation. To test this assumption, Aβ-GFP was overexpressed, its aggregates isolated two days later and analyzed. Mass spectrometry quantification revealed strongly increased co-aggregation of cellular proteins with the Aβ amyloid when the activity of TRMT6/61A was suppressed (Fig. 4a, Table S1 and S2). Based on five biological repetitions, we identified 244 proteins significantly enriched in aggregates in KD cells. There was a considerable overlap between those proteins and the smaller set of 80 co-aggregators from wild-type cells (Fig. 4b). The difference between the sets turned out to be revealing, because the co-aggregome from KD cells contained four functional categories which were not present in the WT set (Fig. 4c). The top-two category was “mRNA binding”, which supported the notion of aberrant sequestration of mRNAs in protein aggregates. In this scenario, amyloid-associated mRNA would attract mRNA binding proteins thus propagating aberrant protein-protein interactions. To verify this inerpretation directly, we used polyT-beads to isolate cellular mRNAs from wild-type and KD cells. Indeed, mRNA association with the amyloid was significantly higher in the KD cells (Fig. 4d). Note the difference of Aβ-GFP accumulation in Fig. 3a (two days after transfection) and Fig. 4d (one day after transfection). We deliberately chose to look for aberrant RNA-protein interactions at an earlier time-point to support their causative relevance for the increased amyloidogenesis.

**Figure 4.**
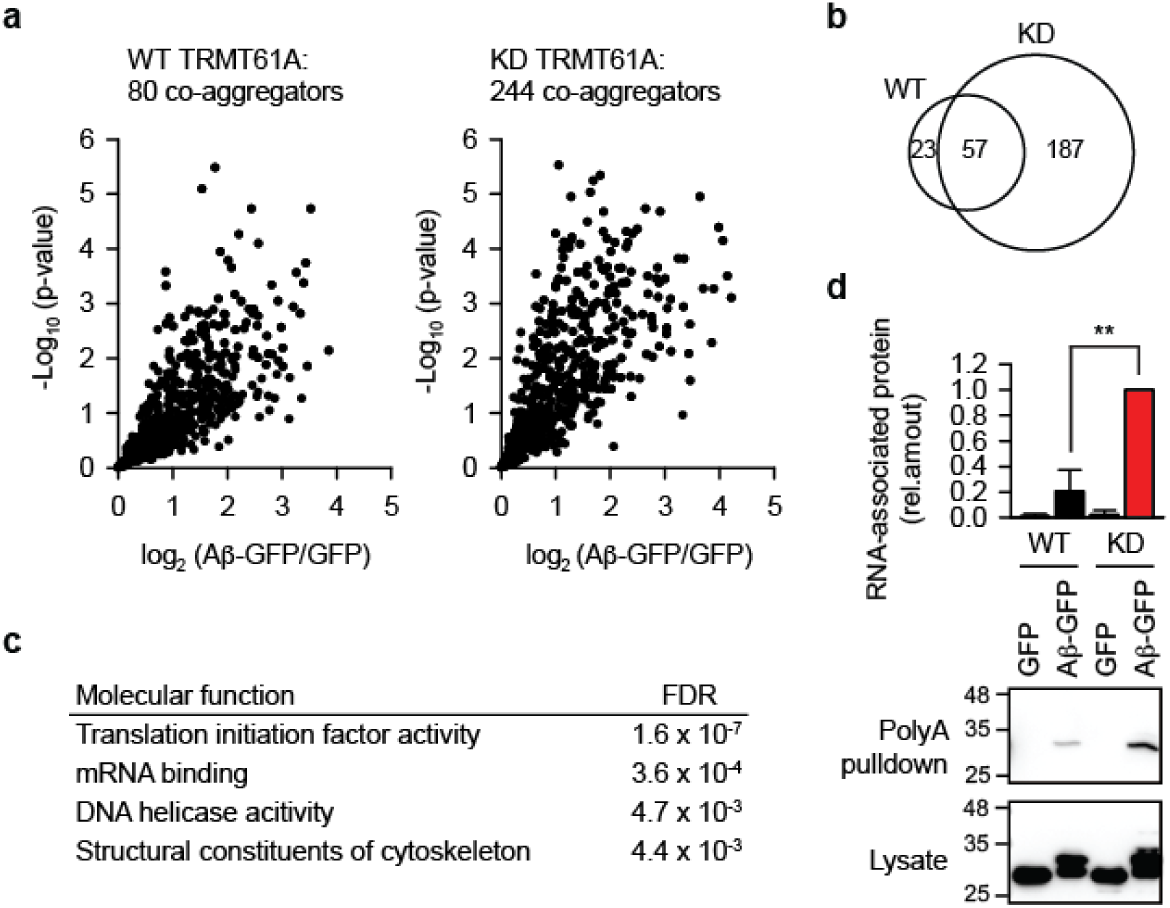
Co-aggregation of proteins and mRNAs is increased under the defective N1-adenine methylation. **a**, Mass spectrometry analysis of proteins that co-aggregate with Aβ-GFP in WT or TRMT61A-deficient HeLa cells 48 hours after transfection. Volcano plot of quantified proteins plotted according to their enrichment on Aβ-GFP over GFP background with the statistical significance of the respective ratios plotted on the y-axis. The size of the determined sets is indicated. N=5 independent experiments. **b**, Overlap between the Aβ-GFP co-aggregators in WT and TRMT61A KD cells. **c**, GO Molecular function categories enriched in the Aβ-GFP co-aggregator set in TRMT61A KD cells, but not in the co-aggregators in WT cells. **d**, Increased amyloid association with mRNAs in TRMT61A KD cells 24 h after Aβ-GFP transfection as determined by polyA pulldowns. Transfection of GFP was used as a control. **p<0.01, two-tailed t-test; N=3 independent experiments (mean + SD).

The enhanced amyloidogenesis and increased mRNA co-aggregation in cells is surprising given the small size of the adenine modification. To explain the far-reaching consequences of the missing small tag, one is tempted to consider the changes of RNA dynamics. It is known that m^1^A blocks base-pairing and induces local melting in RNA molecules (*12*). Together with the fact that the secondary structure of mRNAs is able to determine the specificity of membraneless compartments in cells (*13*), it offers a possible explanation for the aberrant RNA-RNA and RNA-protein associations when TRMT6/61A activity is impaired. mRNAs can be exposed and freed from polysomes following protein synthesis attenuation which takes place upon diverse stressors (*14, 15*). Indeed, our finding of mRNA and Aβ-GFP co-aggregation (Fig. 4d) supports the notion that RNAs lose their native associations not only during heat shock. The enrichment of mRNA binders among the co-aggregators under these conditions suggests a pathogenetic relevance of the aberrant RNA-protein association. RNA repeat expansion disorders provide examples of clinical consequences caused by the RNA binding protein entrapment (*16*). Our study now demonstrates that also normal RNAs can get involved in similar pathogenic loops.

Two questions remain to be answered regarding the model our data propose (Fig. S4). First, the appearance of free mRNA due to the translation impairment under different conditions has to be elucidated. This challenge is especially pressing for the chronic proteostasis stress during amyloidogenesis, where translation factor co-aggregation and protein synthesis defects have been documented (*11, 17*). Second, the extent and mechanism of the m^1^A-tagged mRNA involvement in the protective sequestration of non-methylated mRNAs have to be clarified. The importance of RNA-RNA interactions in sequestration processes have been demonstrated in different experimental settings (*13, 18*).

## Materials and Methods

### Reagents, Plasmids, Antibodies

2,3-Bis(2-methoxy-4-nitro-5-sulfophenyl)-2*H*-tetrazolium-5-carboxanilide inner salt (XTT), phenazine methosulfate (PMS) and cycloheximide were from Sigma-Aldrich, TRIzol and RiboGreen were from Invitrogen. All other chemicals were from Sigma-Aldrich unless otherwise indicated. Aβ-EGFP, Flag-tagged TRMT6 and TRMT61A mammalian expression vectors were purchased from Genscript. Two additional Flag tags where inserted into the vector coding TRMT61A. Luciferase expression vector under T7 promoter was from Promega. Human HSP70 under T7 promoter was cloned into pCH16 vector (a kind gift of H.-C. Chang) for in vitro transcription.

The following antibodies were used: anti-TRMT6 (A303-008A-M) from Bethyl; anti-TRMT61A (sc-107105) from Santa Cruz Biotechnology; anti-eIF2α (9722), anti-eIF2α (phosphor-Ser51) (3398), anti-eIF4E (9742), HRP-conjugated anti-rabbit-IgG (7074) and Alexa Fluor 647-conjugated anti-rabbit IgG (4414) from Cell Signaling; anti-Flag (F1804) and HRP-conjugated anti-mouse IgG (A9044) from Sigma-Aldrich.

### Constructs

Expression constructs generated for this study were prepared by standard molecular biology techniques and coding sequences were verified. Two additional Flag tags where inserted into the vector coding TRMT61A. An m1A motif from human PRUNE1 5’-UTR was introduced into the Ubiquitin-EGFP construct (*9*) to prepare WT-UbE reporter. The motif was mutated such that the to-be methylated adenine was exchanged into uracil to prepare MUT-UbE. Similarly, a 3xFLAG-NQO1 eukaryotic expression vector (*19*) was modified to introduce an m1A motif upstream the coding sequence to construct WT-NQO1 reporter. This motif was mutated as above to prepare MUT-NQO1.

### Cell Lines

The HeLa WT and TRMT61A KD cell lines were cultured in DMEM supplemented with 10% (vol/vol) FBS, 2 mM L-glutamine, 100 IU/mL penicillin G, 100 μg/mL streptomycin sulfate, and nonessential amino acids. HeLa cells with low levels of TRMT61A were prepared from HeLa WT cells by transfection with 1 μg TRM61A Double Nickase Plasmids (h) mix and selection with puromycin.

The B16-F10 cell line was cultured in DMEM supplemented with 10% (vol/vol) FBS, 2 mM L-glutamine, 100 IU/mL penicillin G, 100 μg/mL streptomycin sulfate, and nonessential amino acids.

### Transfections

For Aβ aggregation, 4×10^6^ HeLa cells were electroporated with 30 µg Aβ-GFP or GFP expression vectors and seeded on polylysine-coated cover slides on a 12-well plate at 5×10^5^ cells/well. The medium was refreshed after 6 h. The cells were processed for microscopy or collected and lysed with SDS sample buffer for western blotting 48 h after transfection. For methylation reconstitution, 8×10^6^ HeLa TRMT61A KD cells were transfected by electroporation with 10 µg Aβ-GFP with or without 20 μg Flag-TRMT6 and 20 μg 3xFlag-TRMT61A. The cells were lysed 48 h after transfection. For analysis of the enzymatically impaired TRMT61A mutant D181A, HeLa TRMT61A KD cells were seeded in a 12-well plate at 1×10^5^ cells/well. Next day, cells were cotransfected with 1 µg Aβ-GFP and 1 or 2 µg TRMT61A WT or D181A using polyethylenimine (the ratio DNA:PEI was 1:6 using 1 mg/mL PEI solution). The cells were lysed 24 h after transfection.

### Cellular Viability Assay

HeLa cells were seeded at 1.25×10^4^ cells/well in duplicates in a 48-well plates in 200 μL DMEM supplemented with 25 mM Hepes NaOH pH 7.5. The plates were immediately placed in an incubator at 37°C or 45°C for 2 h and then back to 37°C. 14 h later, medium in each well was discarded and replaced by 200 μL DMEM with 0.33 mg/mL XTT + 12.5 μg/mL PMS (Scudiero et al., 1988). After 2 h of incubation at 37°C, A_475_ and A_600_ were measured. Duplicates were averaged and specific absorbance calculated. To compare sensitivity to arsenite, HeLa WT and TRMT61A KD cells were seeded in two 48-well plates at 1.25×10^4^cells/well in triplicates. Next day, the cells were treated with arsenite for 1 h at 37°C. The cells were washed with PBS, then 200 μL 0.33 mg/mL XTT + 12.5 μg/mL PMS in DMEM was added per well. After 4 h of incubation at 37°C, A_475_ and A_600_ were measured. Triplicates were averaged and specific absorbance was calculated.

### Aβ-GFP Pulldowns for Mass Spectrometry Analysis

8×10^6^ WT or TRMT61A knock-down HeLa cells were transfected by electroporation with 30 µg Aβ-GFP or 10 μg GFP. 48 h after transfection, the cells were washed two times with PBS and resuspended in 300 μL Lysis buffer (10 mM Tris HCl pH 7.4, 100 mM KCl, 5 mM MgCl_2_, 0.5% Deoxycholate, 1% Triton X-100) supplemented with Phosphatase Inhibitor cocktail 2 (1:100), RNasin (1:1000) and Protease inhibitor (1x). The cells were kept on ice for 20 min, then centrifuged at 10.000g at 4°C for 5 min. Supernatants were collected and protein concentration was normalized to 1 μg/μL. 200 μL of each lysate were diluted with 300 μL Dilution buffer (10 mM Tris HCl pH 7.4, 100 mM KCl, 5 mM MgCl_2_) supplemented with RNasin (1:1000) and Protease inhibitor (1x). For each sample, 10 μL GFP-Trap Magnetic Agarose were washed three times with dilution buffer. 450 μL of each lysate were added to 10 μL washed agarose and incubated at 4°C for 1 h with gentle shaking. The agarose was washed three times with Dilution buffer supplemented with RNasin and Protease Inhibitor and three times with 50 mM Tris HCl pH 7.4, 150 mM NaCl. The agarose was frozen at −80°C until further processing for mass spectrometry.

### PolyT Pulldowns of Aβ-GFP

4×10^6^ HeLa cells were electroporated with 30 µg Aβ-GFP or 10 μg GFP expression vectors and seeded in 10 cm dishes (one transfection per dish). 24 h after transfection, the cells were washed twice with ice-cold PBS and resuspended in 300 µL lysis buffer (10 mM Tris HCl pH 7.4, 100 mM KCl, 5 mM MgCl_2_, 0.5% sodium deoxycholate, 1% Triton X-100) supplemented with Phosphatase inhibitor cocktail 2 (1:100) and RNasin (1:1000). After a 15 min incubation on ice, the lysates were centrifuged at 10.000 g for 5 min at 4°C. The protein concentration of the supernatants was measured and normalized. 350 μL lysate were added to 50 µL of washed Oligo d(T)25 Magnetic beads and incubated at RT for 30 min. The beads were washed 3x with lysis buffer and resuspended in 10 μL lysis buffer supplemented with 250 units of Benzonase for elution. After 15 min at 37°C, the supernatant was collected and analyzed by SDS-PAGE and western blotting using anti-GFP antibody.

### WT-UbE and MUT-UbE Reporter Assay

HeLa cells were seeded in a 12-well plate at 1×10^5^ cells per well. Next day, they were transfected with 300 ng reporter constructs using polyethylenimine (PEI) (the ratio DNA:PEI was 1:6 using a 1 mg/mL PEI solution). 24 h after transfection, the cells were treated with 20 μM MG-132 for 3 h, collected and analyzed by western blotting using anti-GFP and anti-GAPDH antibodies.

For the heat shock recovery assay, 12×10^6^ HeLa cells were electroporated with 20 to 30 µg WT-UbE or MUT-UbE and seeded into two 10-cm dishes per transfection. 5 h later, cells were collected, washed and resuspended in serum-free DMEM supplemented with 25 mM Hepes NaOH pH 7.5 and 10 µM Actinomycin D. The cells were heated at 45°C for 30 min and then transferred to 37°C for recovery (timepoint 0). During recovery at timepoints 0, 15 and 30 min, aliquots were collected directly into the pre-heated SDS sample buffer and then analyzed by western blotting using anti-GFP and anti-GAPDH antibodies. To quantify the accumulation of newly synthesized reporter protein, the amount of UbE at timepoint 0 was subtracted.

### WT-NQO1 and MUT-NQO1 Reporter Assay

HeLa cells were seeded in a 12-well plate at 1×10^5^ cells per well. Next day, they were transfected with 250 ng reporter constructs and different amounts of Aβ-GFP construct (0 ng, 250 ng, 500 ng or 1000 ng) using PEI (the ratio DNA:PEI was 1:6 using a 1 mg/mL PEI solution). 24 h after transfection, cells were collected and analyzed by SDS-PAGE and western blotting using anti-GFP and anti-GAPDH antibodies. For quantitative comparison, NQO1 amount per unit of Aβ-GFP was calculated by multiplying the respective NQO1 and Aβ-GFP values. Then, the average for all three Aβ-GFP concentrations was calculated. The average for WT-NQO1 reporter was set as 1.

### Fluorescence Microscopy

For TRMT6/61A and stress granule colocalization analysis, HeLa cells were electroporated with 15 µg each Flag-tagged TRMT6 and TRMT61A and seeded at 0.5×10^6^cells/well in a 12-well plate on polylysine-coated slides. 24 h after transfection, medium was replaced by serum-free DMEM for additional 2 h. The cells were treated with 0.25 mM arsenite for 30 min, washed with PBS and fixed with 3.7% paraformaldehyde/PBS for 9 minutes at RT. They were then permeabilized with acetone at −20°C for 5 min, blocked with 1% BSA/PBS for 1 h at RT and incubated with anti-TIAR in 1% BSA/PBS at RT for 1 h. Incubation with anti-Rabbit IgG-AlexaFluor647 and anti-Flag M2-Cy3 was done in 1% BSA/PBS at RT for 1 h, followed by three washing steps with PBS and DAPI staining. Slides were mounted in PBS and imaged. ImageJ (*20*) was used to compare fluorescence profiles along a set line.

To compare stress granule formation upon arsenite treatment, HeLa WT and TRMT61A KD cells were seeded in a 12-well plate with polylysine-coated cover slides at 1.5×10^5^cells/well. Next day, medium was replaced by serum-free medium and the cells were kept at 37°C for 2 h. Arsenite was then added at 0.0625 mM for 30 min at 37°C. Cells were washed with PBS, fixed with 3.7% paraformaldehyde at RT for 10 min, permeabilized with acetone at −20°C for 5 min, blocked with 1% BSA/PBS for 1 h at RT and incubated with anti-TIAR for 1 h. Incubation with anti-Rabbit-IgG Alexa Fluor647 conjugate for 1 h, three washes with PBS and staining with DAPI followed. Slides were imaged using a Zeiss LSM-780 inverted confocal microscope with a 63x oil immersion objective. The experiment was performed in triplicates and at least 200 cells were analyzed per condition and per repetition. Quantification was performed using CellProfiler by detecting nuclei and identifying by propagation from the nuclei. SGs were identified and counted within each cell.

For aggregation analysis, HeLa cells transfected with Aβ-GFP or control GFP were fixed 48 h after transfection with 4% paraformaldehyde/PBS at RT for 1 h and stained with DAPI. Cells were imaged using a Zeiss LSM-780 inverted confocal microscope with a 40x oil immersion objective. Using CellProfiler, nuclei were detected and cells identified by propagation from the nuclei. Fluorescence intensity was measured for each cell. Dead or dying cells were filtered out based on intensity values. The experiment was performed in triplicates and at least 200 GFP-positive cells were analyzed per condition and per repetition to determine the fraction of cells with aggregates.

### Quantitative PCR

Total RNA was extracted with TRIzol. cDNAs were prepared using RevertAid kit and diluted 10x. Reporter and GAPDH sequences were amplified using respective primers and KiCqStart SYBR Green qPCR ReadyMix. The cycling conditions used were the following: 3 min at 95°C, 39x (15 sec 95°C, 30 sec 58°C, 15 sec 72°C). Each reaction was performed in triplicates and results were analyzed as described (*21*) with GAPDH as a reference. Analysis was performed using the CFX Manager Software (Bio-Rad).

### tRNA Isolation for HPLC Analysis

RNAs shorter than 200 nucleotides were extracted using mirVana miRNA Isolation kit. The extracted RNAs were separated in an 8% Polyacrylamide-TBE-urea gel, the tRNA band was visualized by UV shadowing and cut out of the gel. The gel pieces were incubated overnight with 0,5 M ammonium acetate at 25°C and 600 rpm. Next day, the gel pieces were discarded and 1 mL 100% ethanol was added per 400 μL of extract. The mixture was incubated at −80°C for 1 hour and tRNAs were pelleted at 17.000 g for 30 minutes at 4°C. Pellets were washed with 70% ethanol, air-dried and frozen until further processing.

### HPLC Analyses

tRNA pellets were resuspended with 16 µL of MS grade water and digested to nucleosides by incubation with 0.1 U nuclease P1 and 0.1 U bacterial alkaline phosphatase BAP C75 in P1 buffer (20 mM NH_4_OAc pH 5.3) at 37 °C for 1 h. Chromatographic separation was performed on an UltiMate 3000 HPLC (Thermo Fisher Scientific) equipped with a diode array detector (DAD). 5 µL of the nucleoside mixture were loaded on a Synergy Fusion RP column (4 mm particle size, 80Å pore size, 250 mm length, 2 mm inner diameter) from Phenomenex (Aschaffenburg, Germany) with 100% buffer A (5 mM NH_4_OAc pH 5.3) and separated at 35°C with a flow rate of 0.35 ml/min using a linear gradient of buffer B (100% acetonitril) to 20% in 20 min, then to 40% in 2 min. At the end of the gradient the column was washed with 95% buffer B for 7 min and re-equilibrated with buffer A for 4 min. The nucleosides were monitored by recording the UV chromatogram at 254 nm.

### Quantification of Modified Nucleotides by Mass Spectrometry

Stress granules were prepared as described (*6*) omitting the affinity purification step. HeLa cells were treated with 0.5 mM Arsenite at 37°C for 60 min, washed, pelleted and flash-frozen in liquid nitrogen. The pellets were thawed, resuspended in 500 µL lysis buffer (50 mM Tris HCl pH 7.4, 100 mM potassium acetate, 2 mM magnesium acetate, 0.5 mM DTT, 50 µg/mL Heparin, 0.5% NP40, Protease inhibitor and RNasin), passed seven times through a 26 G needle and centrifuged at 1000 g for 5 min at 4°C. The lysate was transferred to a new tube and centrifuged at 17000 g for 20 min at 4°C. The resulting supernatant was called ‘Soluble 1’. The pellet was resuspended in 500 µL lysis buffer and centrifuged again at 17000 g for 20 min at 4°C. The resulting supernatant was called ‘Soluble 2’. The pellet was resuspended in 150 µL lysis buffer, spinned at 850 g for 2 min at 4°C. The supernatant was called ‘SG pellet’ and constituted the stress granule core enriched fraction. Total RNA was extracted from the SG pellet by Trizol extraction and RNA precipitation and stored at −80°C until MS analysis.

PolyA mRNA for comparison was prepared from HeLa cells heated at 45°C for 60 min. The cells were lysed in 500 µL lysis buffer (20 mM Tris HCl pH 7.5, 500 mM LiCl, 0,5% LiDS, 1 mM EDTA, 5 mM DTT), incubated at RT for 5 min and passed 5x through a 20 G needle. The lysate was added to 100 μL of washed Oligo-d(T)25 Magnetic beads and incubated at RT for 10 min. The beads were washed 2x with WB-I (20 mM Tris HCl pH 7.5, 500 mM LiCl, 0,1% LiDS, 1 mM EDTA, 5 mM DTT), 2x with WB-II (20 mM Tris-HCl pH 7.5, 500 mM LiCl, 1 mM EDTA) and 1x with Low Salt Buffer (20 mM Tris HCl pH 7.5, 200 mM LiCl, 1 mM EDTA). The beads were resuspended in 100 µL Elution buffer (20 mM Tris HCl pH 7.5, 1 mM EDTA), mRNAs were eluted by heating at 50°C for 2 min, precipitated and stored at −80°C until MS analysis.

RNA from stress granules or cytosolic mRNAs from heated HeLa cells were resuspended in 10 μl of MS water and digested with 0.05 units of nuclease P1 (Wako) and 0.05 units of bacterial alkaline phosphatase (BAP C75, Takara) in P1 buffer (20 mM ammonium acetate pH 5.3) for 1 h at 37°C. Samples were store at −20°C before mass spectrometry.

Chromatographic separation of the nucleosides was performed with an Easy-nLC1000 on an in-house packed column (100 µm inner diameter, 50 cm length, 4 μm Synergi Fusion 80 Å pore size from Phenomenex (Aschaffenburg, Germany) using a gradient from mobile phase A (5 mM ammonium formate pH 5.3) to 32% mobile phase B (100% acetonitrile) for 50 min followed by a second step to 40% B for 8 min, with a flow rate of 200 nl/min.

Retention times were monitored using 10 nM solutions of nucleoside standards purchased from Sigma Aldrich (main nucleosides) and Carbosynh (N1-methyladenosine and N6-methyladenosine). Nucleosides were injected into a Q Exactive Plus mass spectrometer equipped with a Nanospray Flex Ion-Source and analysed in the positive mode with a method consisting of full MS and targeted SIM scans. Full scans were acquired with AGC target value of 10^6^, resolution of 70.000 and maximum injection time of 100 ms. Adenosine, N1-methyladenosine and N6-methyladenosine were monitored at the respective retention time with a 4-amu isolation window, AGC target value of 2×10^5^, 140.000 resolution and maximum injection time of 500 ms.

The correct identification of N1- and N6-methyladenosines was proved by fragmentation of the parent ion using a normalized collision energy of 30 and recording MS/MS scans at a resolution of 70000, 4.0 m/z isolation window, 2×10^5^ ACG target and maximum fill time of 120 ms.

Manual identification and quantification of nucleosides was performed using Thermo Xcalibur Qual Browser. Extracted ion chromatograms were generated with a deviation of 0.002 Da from the theoretical m/z value and the peaks were integrated using the manual peak annotation function. Absolute amounts of nucleosides were calculated generating an external calibration curve in the range 0.2-3000 fmol by serial dilutions of the nucleoside standards. N1-methyladenosine (m^1^A) and N6-methyladenosine (m^6^A) amounts were reported as percentage of adenosine (A).

### Quantitative Mass Spectrometry

#### Sample preparation

Pulled down proteins were processed on-beads for LC-MS/MS analysis as following. Beads were re-suspended in 50 µL 8M urea/50 mM Tris HCl pH 8.5, reduced with 10 mM DTT for 30 min and alkylated with 40 mM chloroacetamide for 20 min at 22°C. Urea was diluted to a final concentration of 2 M with 25 mM Tris HCl pH 8.5, 10% acetonitrile and proteins were digested with trypsin/lys-C mix overnight at 22°C. Acidified peptides (0.1% trifluoroacetic acid) were desalted and fractionated on combined C18/SCX stage tips (3 fractions). Peptides were dried and resolved in 1% acetonitrile, 0.1% formic acid.

#### LC-MS/MS

LC-MS/MS was performed on a Q Exactive Plus equipped with an ultra-high pressure liquid chromatography unit (Easy-nLC1000) and a Nanospray Flex Ion-Source (all three from Thermo Fisher Scientific). Peptides were separated on an in-house packed column (100 μm inner diameter, 30 cm length, 2.4 μm Reprosil-Pur C18 resin) using a gradient from mobile phase A (4% acetonitrile, 0.1% formic acid) to 30% mobile phase B (80% acetonitrile, 0.1% formic acid) for 60 min followed by a second step to 60% B for 30 min, with a flow rate of 300 nl/min. MS data were recorded in data-dependent mode selecting the 10 most abundant precursor ions for HCD with a normalized collision energy of 30. The full MS scan range was set from 300 to 2000 m/z with a resolution of 70000. Ions with a charge ≥2 were selected for MS/MS scan with a resolution of 17500 and an isolation window of 2 m/z. The maximum ion injection time for the survey scan and the MS/MS scans was 120 ms, and the ion target values were set to 3×10^6^ and 10^5^, respectively. Dynamic exclusion of selected ions was set to 30 s. Data were acquired using Xcalibur software.

#### Data analysis with MaxQuant

MS raw files from five biological replicates of pulldown and background samples were analyzed with Max Quant (version 1.5.3.30) (Cox and Mann, 2008) using default parameters. Enzyme specificity was set to trypsin and lysC and a maximum of 2 missed cleavages were allowed. A minimal peptide length of 7 amino acids was required. Carbamidomethylcysteine was set as a fixed modification, while N-terminal acetylation and methionine oxidation were set as variable modifications. The spectra were searched against the UniProtKB human FASTA database (downloaded in November 2015, 70075 entries) for protein identification with a false discovery rate of 1%. Unidentified features were matched between runs in a time window of 2 min. In the case of identified peptides that were shared between two or more proteins, these were combined and reported in protein group. Hits in three categories (false positives, only identified by site, and known contaminants) were excluded from further analysis. For label-free quantification (LFQ), the minimum ratio count was set to 1.

#### Data analysis with Perseus

Bioinformatic data analysis was performed using Perseus (version 1.5.2.6) (Tyanova et al., 2016). Proteins identified in the pulldown experiments were further included in the analysis if they were quantified in at least 4 out of 5 biological replicates in at least one group (pulldown/background). Missing LFQ values were imputed on the basis of normal distribution with a width of 0.3 and a downshift of 1.3. Proteins enriched in the pulldown were identified by two-sample t-test at a permutation-based FDR cutoff of 0.05 and s0 = 0.1. Categorical annotations were added in Perseus and a Fisher’s exact test with a p-value threshold of 0.001 was run for GO term enrichment analysis.

### m1A Motif

The reference set of human genes (11,195 genes) was taken from Khong et al (Khong et al., 2017). Ensemble Biomart (version 90) was used to select all transcript variants (69,090 transcripts) and to retrieve the cDNA sequences. The unique transcripts were filtered to select those which match the exact length reported in Khong et al (Khong et al., 2017) (final set of 9,301 transcripts). The motifs centered on TTCAA (AGTTCAANNCT, CGTTCAANNCG, GGTTCAANNCC, TGTTCAANNCA) and centered on TTCGA (AGTTCGANNCT, CGTTCGANNCG, GGTTCGANNCC, TGTTCGANNCA) were searched separately in all the sequences. A search algorithm was written in Python for this specific task.

### Bioinformatics Analysis

GO terms enrichment analysis of mRNAs with m1A motif enriched in SG (Khong et al., 2017) and filtered as described above was performed on line with Panther (Mi et al., 2017) using the Overrepresentation Test tool (released 2017-12-05) and the GO ontology database (released 2017-11-28). A binomial test with p-value threshold of 0.05 was run using the set of mRNAs neither enriched nor depleted from SG as reference. GOslim Molecular Function enrichment analysis of the Aβ-GFP coaggregome was performed with Panther as described above. As input gene names of Aβ-GFP interactors and of the human genome (all genes in database) were used. FDR cut off for the Fisher’s exact test was set to 0.05.

### Statistical analyses

All repetitions in this study were independent biological repetitions performed at least three times if not specified differently. To identify significantly increased proteins in pulldowns (RNA over background binding) in mass spectrometry analyses, a two-sample t-test analysis of grouped biological replicates was performed using a FDR cutoff of 0.05 with s0 = 0.1. Categorical annotation was added in Perseus and a Fisher exact test with a p-value threshold of 0.05 was run for GOslim term enrichment analysis. Statistical significance for categorical readouts in microscopy was analysed by chi-square analysis. Means and standard deviations were calculated from at least three independent experiments.

## Acknowledgments

We thank H. Schwalbe for critical discussion. We thank the ERC (StG-311522 to R.M.V., StG-309545 to G.G.T) and DFG (EXC115 to R.M.V) for funding. M.H. is funded by DFG CRC „Molecular Mechanisms of RNA-mediated Regulation”.

## Author Contributions

R.M.V. conceived and supervised the project. R.M.V., M.H. and G.G.T. conceived the experiments. M.A., A.M.-L. and G.H. designed and performed the experiments and analyzed the data. G.C. performed mass spectrometry and analyzed the data. R.D.P., G.G.T. and G.C. performed bioinformatics analyses. R.M.V. wrote the manuscript with contribution from all authors.

## Competing Financial Interests

The authors declare no competing financial interests.

## Supplementary Materials

Table S1. MaxLFQ Quantitative Data and Identifiers of Aβ-GFP Interactors

Table S2. Aβ-GFP Interactors in WT and TRMT61A knock-down HeLa cells

**Figure S1.**
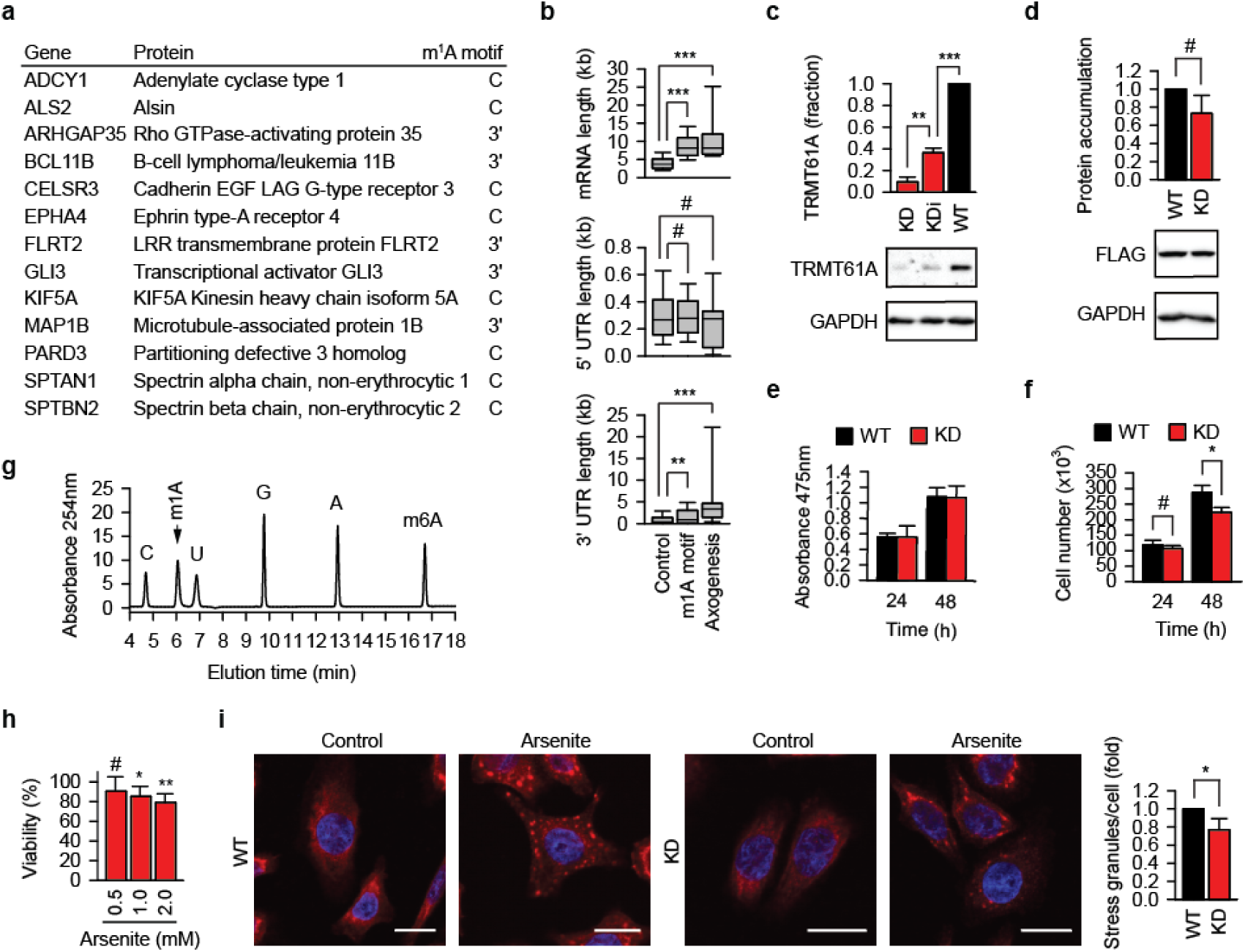
Characterization of TRMT6/61A and its motif during acute proteostasis stress. **a**, The list of human genes from GO category “Axonogenesis” that encode mRNA which are enriched in SG and contain an m^1^A motif. The position of the motif in the transcript is indicated (C, coding region; 3’, 3’-UTR). **b**, Distribution among the indicated sets of mRNAs of total, 5’-UTR and 3’-UTR length. Control, mRNA set neither enriched nor depleted in SG; m1A motif, mRNAs enriched in SG and containing the m^1^A motif; Axonogenesis, SG-enriched mRNA with the m^1^A motif which belong to the GO category “Axonogenesis”. **p<0.01, ***p<0.001, Mann-Whitney test. #, not significant difference. **c**, TRMT61A amount determined by western blotting. GAPDH was used as loading control. WT, wild-type HeLa cells; KDi, knock-down with intermediate level of TRMT61A; KD, knock-down with strong reduction of TRMT61A level. **p<0.01, ***p<0.001, two-tailed t-test; N=3 independent experiments (mean + SD). **d**, Protein synthesis is not affected significantly in TRMT61A knock-down cells as exemplified by the accumulation of Flag-NQO1 during 24 h after transfection. GAPDH was used as loading control. WT, wild-type HeLa cells; KD, knock-downs of TRMT61A. #, not significant difference, two-tailed t-test; N=3 independent experiments (mean + SD). **e**, Cell viability at 37°C is not affected significantly by the reduced level of TRMT61A as measured by XTT assay. WT, wild-type HeLa cells; KD, knock-downs of TRMT61A. #, not significant difference, two-tailed t-test; N=3 independent experiments (mean + SD). **f**, Cell proliferation at 37°C is not affected significantly by the reduced level of TRMT61A. WT, wild-type HeLa cells; KD, knock-downs of TRMT61A. #, not significant difference, two-tailed t-test; N=3 independent experiments (mean + SD). **g**, HPLC analysis of the 10 μM mixture of the ribonucleosides cytidine (C), N1-methylated adenosine (m^1^A), uridine (U), guanosine (G), adenosine (A) and N6-methylated adenosine (m6A). **h**, Viability of TRMT61A knock-down cells after arsenite treatment for 1 h in comparison (%) to wild-type cells at the same concentrations of arsenite. *p<0.05, **p<0.01, two-tailed t-test; #, not significant difference. N=3 independent experiments (mean + SD). **i**, Immunofluorescence staining of TIAR (red) to detect stress granule formation upon arsenite treatment for 30 min in serum-free medium. DAPI staining (blue), nuclei. Scale bar 20 μm. WT, wild-type HeLa cells; KD, knock-downs of TRMT61A. *p<0.05, two-tailed t-test; N=3 independent experiments (mean + SD).

**Figure S2.**
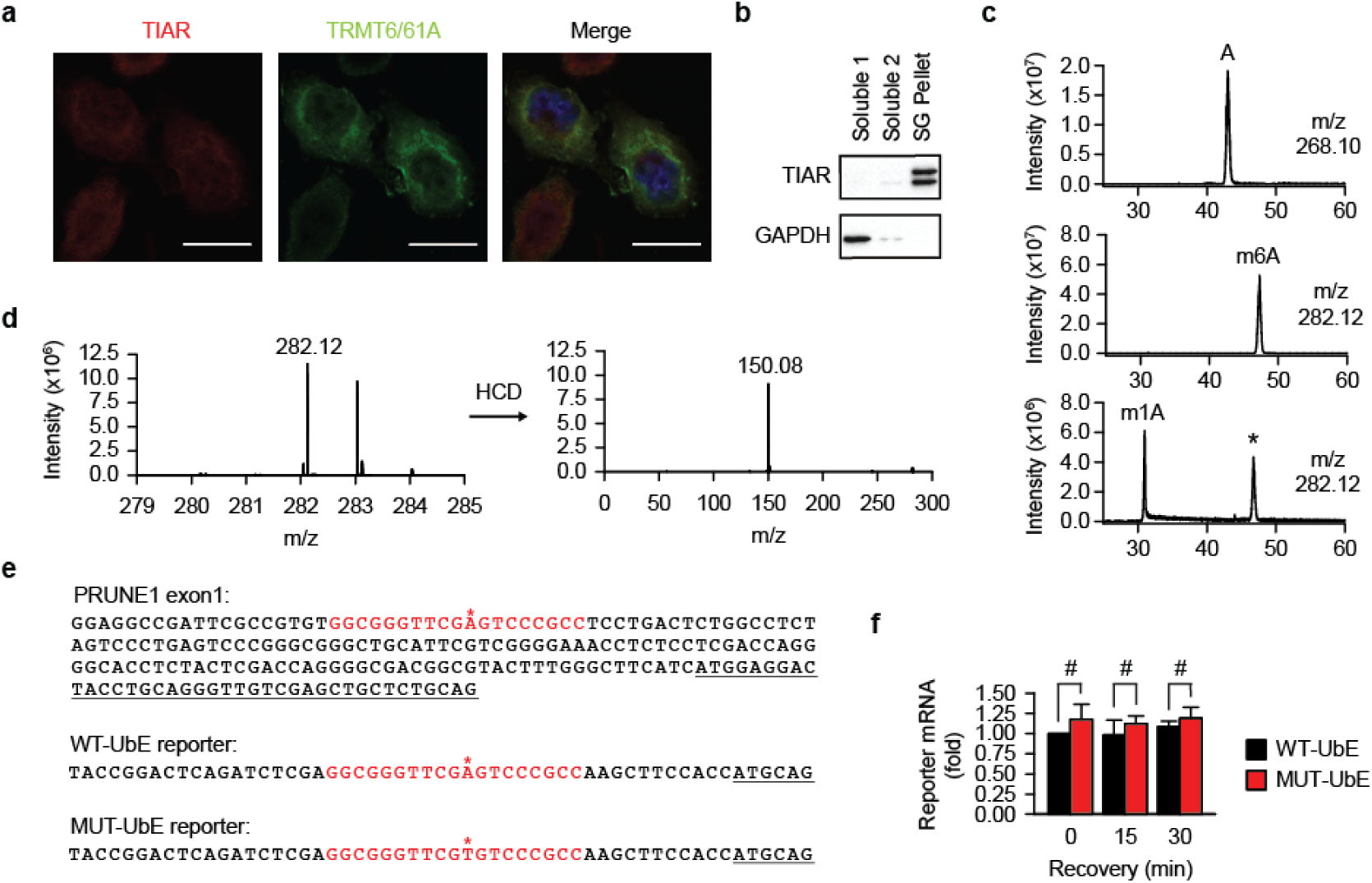
Experimental tools to analyze mRNA methylation during heat shock. **a**, Localization of TRMT6/61A and TIAR in control cells. A representative image from three independent experiments. Scale bar 20 μm. Red, TIAR; green, TRMT6/61A. **b**, A representative western blot of stress granules (SG pellet) used for mass spectrometry of methylated adenosine (N=3 independent experiments). GAPDH was used as cytosolic marker. Soluble 1, Soluble 2, supernatant fractions as described in Method details. **c**, Extracted ion chromatograms of indicated m/z with a range of ±0.002 Da. Top, a mixture of four standard ribonucleosides plus N1-methyladenosine (m1A) and N6-methyladenosine (m6A); middle, N6-methyadenosine only; bottom, N1-methyladenosine only. A, adenosine. *, Dimroth rearrangement of m^1^A to m^6^A. **d**, Spectra of the parent ion (m/z = 282.12) and fragment ion (m/z = 150.08) upon higher-energy collisional dissociation (HCD). Selected ion monitoring was performed at m/z = 282.12 ±4 Da. **e**, m^1^A-motif in the 5’-untranslated region of the PRUNE1 gene and in the WT-UbE and MUT-UbE reporters is marked red. Asterisk indicates adenine methylated by TRMT6/61A. Translated sequence is underlined. **f**, The amounts of reporter mRNA during recovery after heat shock do not differ significantly as determined by quantitative PCR. #, not significant difference, two-tailed t-test; N=3 independent experiments (mean + SD).

**Figure S3.**
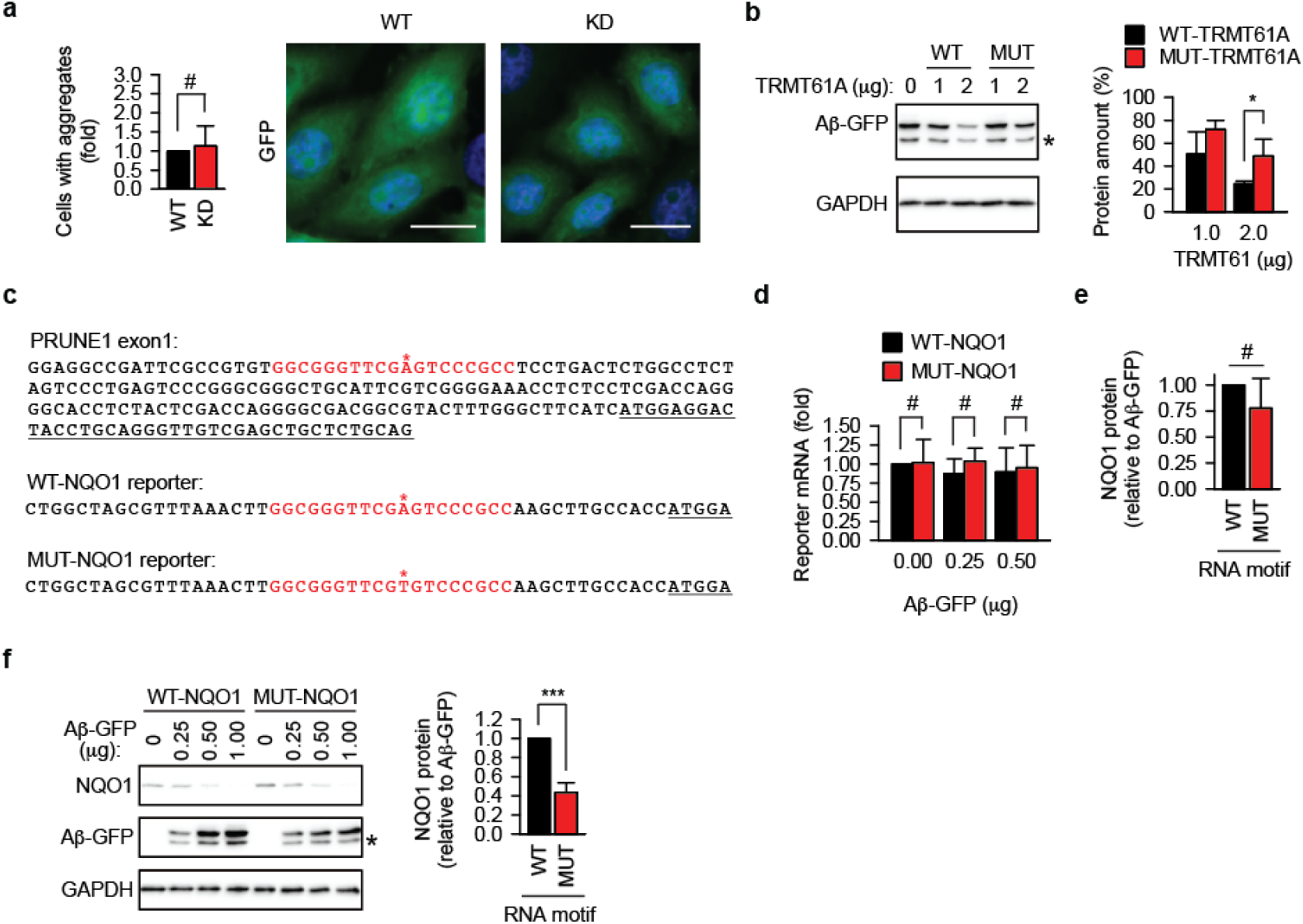
Analysis of amyloidogenesis in RNA methylation-insufficient cells. **a**, GFP analysis in transiently transfected wild-type (WT) and TRMT61A knock-down (KD) cells. Scale bar 20 μm. #, not significant difference, chi-square test on cumulative data from three independent experiments (N=3, mean + SD). **b**, Activity-impaired TRMT61A mutant (MUT-TRMT61A) cannot suppress the accumulation of Aβ-GFP as efficiently as the wild-type enzyme (WT-TRMT61A) in TRMT61A knock-down HeLa cells. Cells were analyzed by western blotting 24 h after transfection. GAPDH was used as loading control. *p<0.05, two-tailed t-test (N=4, mean + SD). **c**, m^1^A-motif in the 5’-untranslated region of the WT-NQO1 and MUT-NQO1 reporters is marked red. Asterisk indicates adenine methylated by TRMT6/61A. Translated sequence is underlined. **d**, The amounts of reporter mRNA do not differ significantly upon 24 h coexpression with Aβ-GFP as determined by quantitative PCR. #, not significant difference, two-tailed t-test; N=3 independent experiments (mean + SD). **e**, The difference of NQO1 protein accumulation from WT-NQO1 (WT) and MUT-NQO1 (MUT) reporter in the presence of Aβ-GFP is not significant in TRMT61A knock-down HeLa cells. #, not significant difference, two-tailed t-test; N=3 independent experiments (mean + SD). **f**, Accumulation of NQO1 protein from WT-NQO1 and MUT-NQO1 transcripts 24 h after cotransfection with indicated amounts of Aβ-GFP in murine melanoma B16-F10 cells. One representative anti-FLAG western blot (NQO1) is shown. GAPDH was used as loading control. For quantitative comparison, NQO1 amount per unit Aβ-GFP was calculated and averaged for all three Aβ-GFP concentrations. This value for WT-NQO1 reporter was set as 1. ***p<0.001, two-tailed t-test; N=4 independent experiments (mean + SD).

**Figure S4.**
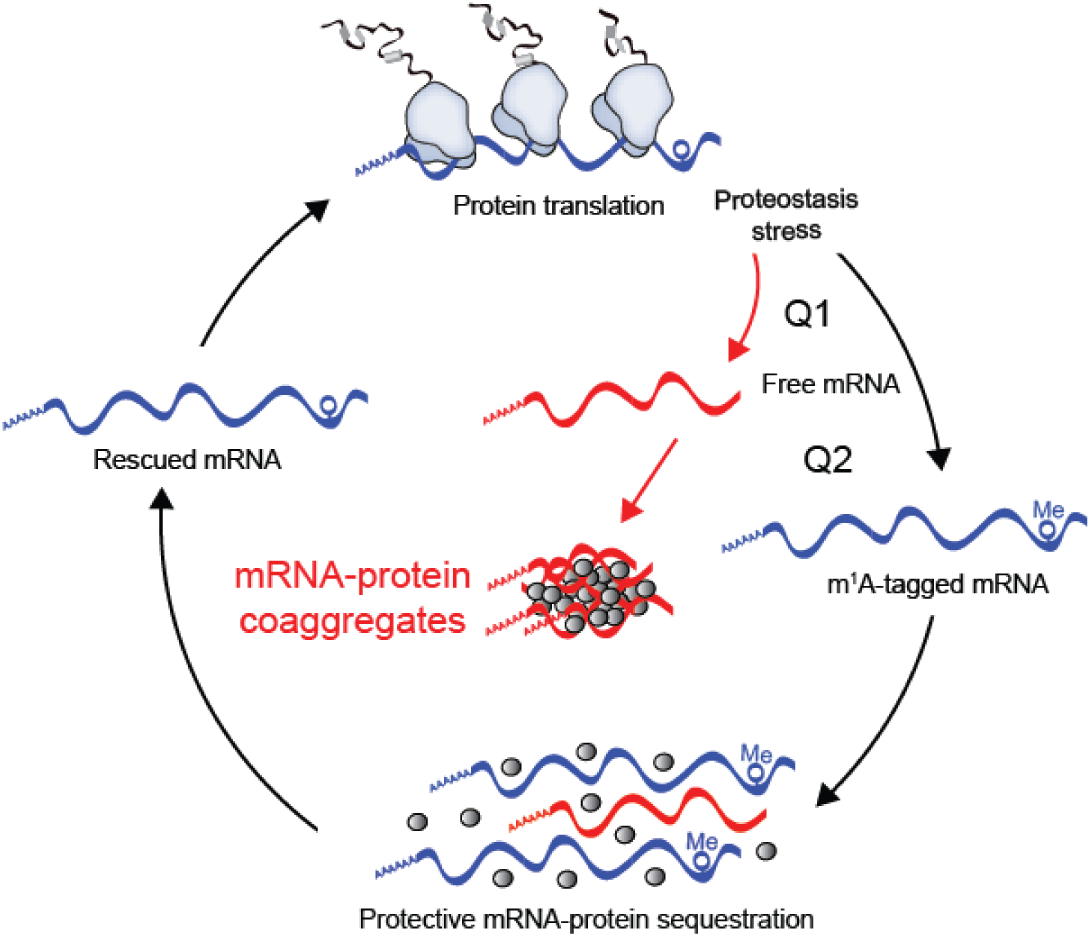
Orderly mRNA sequestration and aberrant RNA-protein association during proteostasis stress. A model of mRNA and protein homeostasis during acute (heat shock) and chronic (amyloidogenesis) proteostasis stress. Two main questions remain open. Q1: Molecular details of the free mRNA appearance during different types of stress. Q2: The extent and mechanism of m^1^A-tagged mRNA (blue) involvement in the protective sequestration of non-methylated mRNA (red).

